# Canada goose fecal microbiota correlate with geography more than host-associated co-factors

**DOI:** 10.1101/2022.07.07.499127

**Authors:** Joshua C. Gil, Celeste Cuellar, Sarah M. Hird

## Abstract

The gut microbiota has many positive effects on the host, but how the microbiota is shaped and influenced can vary greatly. These factors affect the composition, diversity, and function of host-associated microbiota; however, these factors vary greatly from organism to organism and clade to clade. The avian microbiota often correlates more with the sampling locations rather than host-associated co-factors. These correlations between location and microbiota often only include a few sampling locations within the species’ range. To better understand the connection between geographic distance and the microbiota, were collected from non-migratory Canada geese across the United States. We expected host-associated factors to have minimal effect on the microbiota and geese microbiota will be strongly correlated to geography. We hypothesized more proximal geese will be exposed to more similar environmental microbes and will have more similar microbiota. Canada geese microbiota are largely similar across the entire sampling range. Several bacterial taxa were shared by more than half of the geese. Four phyla were found in the majority of the samples: *Firmicutes, Proteobacteria, Bacteroidetes*, and *Actinobacteria*. Three genera were also present in the majority of the samples: *Helicobacter, Subdoligranulum, and Faecalibacterium*. There were minimal differences in alpha diversity with respect to age, sex, and flyway. There were significant correlations between geography and beta diversity. Supervised machine learning models were able to predict the location of a fecal sample based on taxonomic data alone. Distance decay analysis show a positive relationship between geographic distance and beta diversity. Our work provides novel insights into the microbiota of the ubiquitous Canada goose and further supports the claim that the avian microbiota is largely dominated by the host’s environment. This work also suggests that there is a minimum distance that must be reached before significant differences in the microbiota between two individuals can be observed.

## INTRODUCTION

All animals interact with microorganisms and many form symbiotic associations with their microbiota (the collection of all microorganisms living in a particular environment) [1]. Microbes can confer numerous benefits to their hosts by promoting normal immune function [2–4], pathogen exclusion [5,6], and nutritional aid [7–10] promoting host fitness and health. The host, in return, provides a stable environment and available nutrients [11]. Most microbiota research has been conducted on mammals and avian microbiota research typically focuses on economically important species [12]. Comparatively little is known about the non-poultry bird microbiota even though they are a diverse and globally important lineage with over 10,000 extant species [13]. Growing evidence shows many of the patterns in mammalian microbiota do not apply to birds [14]. Birds vary in migratory habits, diet, flight patterns, behavior, mating strategies, longevity, etc., and all of these factors may impact the microbiota [15]. There is substantial evidence that avian microbiota are often not strongly associated with their host species [16, 17], rather their microbiota exhibit great intraspecific variation with emphasis on location, age, and diet [12,13,17–24]. This relatively weak relationship between host phylogeny and gut microbiota diversity may be a consequence of multiple, non-exclusive factors: variable diets between species and throughout the year, frequent and early introduction of environmental microbes (e.g., from the nest), or the various ways that selection for powered flight has affected bird anatomy and physiology [11].

Extrinsic and intrinsic factors can affect the microbiota to varying degrees. Extrinsic factors originate from outside of the host and include the host’s environment [12]. Birds are directly exposed to potential gut microbes in their preferred habitats, including vegetation, nesting materials, water, and soil. The microbes in the gut will also be affected by the types of food the host eats and the availability of different food types in the local environment. Therefore, birds from closer geographic locations should be exposed to similar environmental pressures, food sources, and microbes culminating in a more similar microbiota compared to birds that are further apart [15], although it is unclear when the differences in the environment will result in a difference in the microbiota. Intrinsic factors originate from within the host, such as genetics, age, sex, and health. Host taxonomy can strongly correlate to bird microbiota [15, 19]. In addition, males and females have different physiological traits and behaviors which can influence their microbiota composition. The importance of intrinsic factors and extrinsic factors varies from species to species [20,25,26]. Understanding the variation within a species across its entire natural range is important to put population or site level variation of the microbiota in the proper context yet can be logistically challenging. Herein, we aimed to quantify the relationship between the microbiota and the geographic distribution of its host species. Geography is an important co-variable in microbial community composition, where microbial communities that are closer geographically share more characteristics; whereas, more distant communities have more divergent microbial communities [27, 28]. This general pattern makes intuitive sense, as myriad biotic and abiotic variables are correlated to geographic space, but the shape of this relationship within host-associated microbiota is poorly known. We hypothesized that the Canada geese microbiota would follow this same trend, with birds from the same site having the most similar microbiota and birds from opposite sides of the country having the most different. Alternatively, it may be that the host intrinsic pressures are so strong that the microbiota would be relatively consistent across the entire population.

Canada geese (*Branta canadensis*) are thriving in North America, have disparate migratory behaviors, and are of human interest due to the risk of their pathogen-containing feces contaminating water reservoirs [29].They are also known to be carriers of pathogenic bacteria of *Campylobacter spp*. and *Salmonella spp*. posing possible threats to public health [30]. The Canada goose diet shifts depending on the season, age of the bird, and resource availability [31]. Canada geese have a large range across the United States; however, they are predominantly folivores and have the same general diet, primarily grasses, regardless of their location [32]. This reduces the noise introduced by a diverse diet and will highlight the signal generated by their geographic distribution.

In this study, we characterized the gut microbiota of non-migratory Canada goose populations from eight states across the United States, representing three major waterfowl flyways (migratory paths that ducks and geese follow every year going south to winter and north to breed). We collaborated with state Fish and Wildlife Departments and obtained fecal samples from the Pacific (Washington and Nevada), Central (Utah and Kansas), and Atlantic (New Jersey, New York, Connecticut, Rhode Island) flyways in the summer of 2019, when geese were molting (flightless). We hypothesize that Canada goose populations that are geographically closer together will be exposed to more similar environmental microbes and therefore, will possess more similar gut microbiota than geese from more distant populations. We expect to see this manifest in more similar microbiota within a state and within a flyway.

## METHODS

### Sample collection and preservation

Fecal sample collection was done during early summer 2019 in collaboration with Fish and Wildlife agencies from nine states: Washington (WA), Oregon (OR), California (CA), Nevada (NV), Kansas (KS), New Jersey (NJ), New York (NY), Connecticut (CT), and Rhode Island (RI). WA, CA, OR, NV, KS, and NJ were sent sample collection kits containing 30-100 freezer tubes filled with 700 uL of ZymoBionics DNA/RNA shield (Zymo Research, Irvine, California, USA) to preserve DNA at room temperature, sterile single-use tweezers, pen and notebook to record pertinent data. Samples were also collected in NY, CT, and RI with the same sample collection kits. All samples were collected when the Canada geese were molting their flight feathers and were unable to fly. This limited the potential effects of seasonal variations as they molt in early summer. Geese were being handled for annual banding and population census counts. Fecal samples were collected passively during banding using the sterile tweezers within five minutes of deposition and only feces that was not in contact with the ground were collected to avoid contamination from the environment. Approximately 70 uL of the feces was placed into the 700 uL of DNA/RNA shield maintaining the 10% volume ratio for maximum preservation. The location and band number were recorded for every sample. In some states, the sex and age (Adult: >1 year old and Juvenile: <1 year old) of the bird were recorded as well. After collection each day, samples were stored in a 4°C fridge or −10°C freezer until the samples could be shipped to the University of Connecticut and stored in a −80°C freezer until extraction. In total,419 samples were collected (Table 1).

**Table 1:**
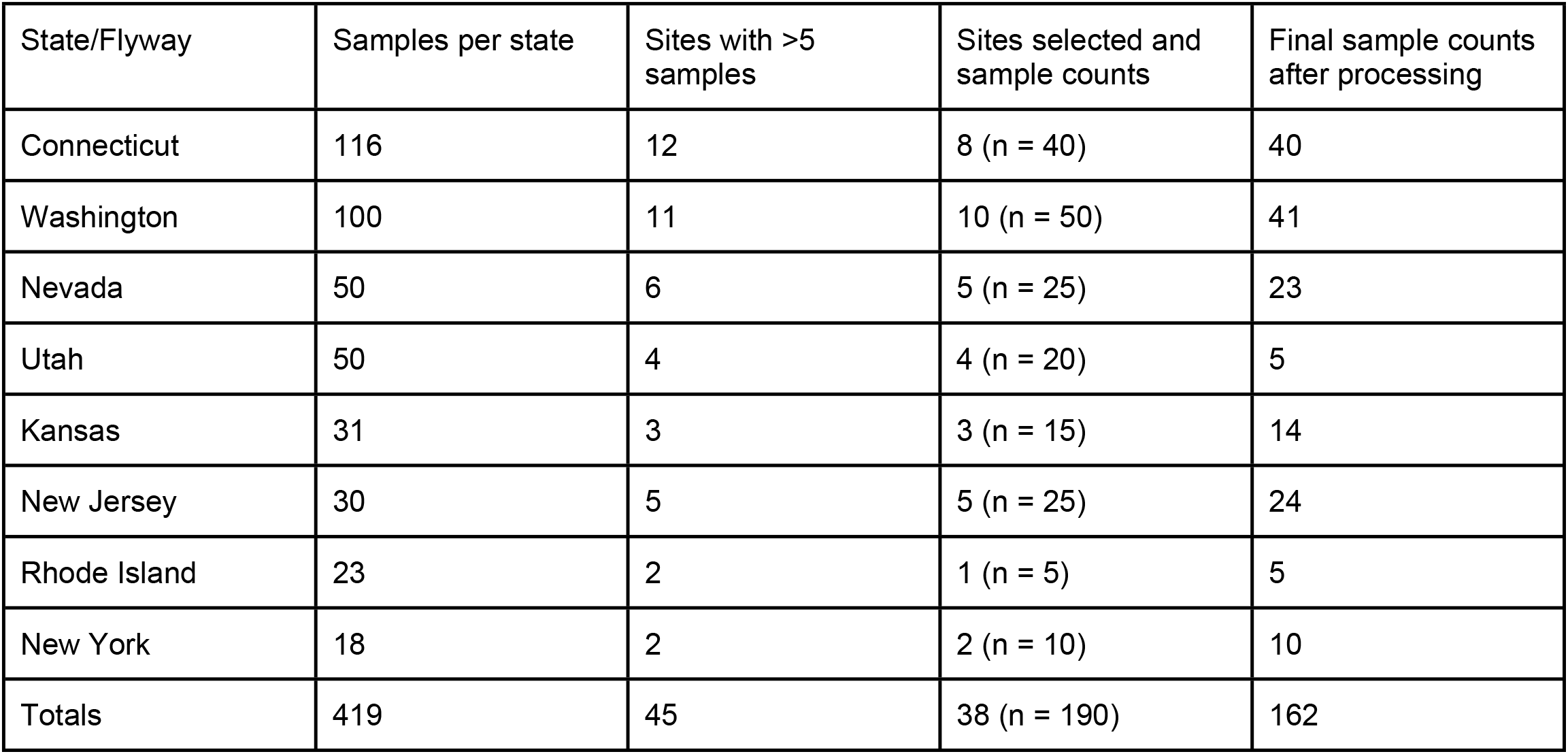
Sample counts and locations.

### Nucleic Acid Extraction

38 sampling locations, that each had at least five samples, were selected for analysis. The 190 samples were extracted in the summer of 2021 using ZymoBionics DNA miniprep Kit (Zymo Research, Irvine, California, USA). We used the standard extraction protocol with modifications: the Lysis buffer provided was not used and instead, an additional ZymoBionics DNA/RNA shield was used as a substitute, the bead beating step was increased to 30 minutes, and a double final elution using 75 uL of 60°C DNase/RNase free water. Negative extraction controls were included for every 25 samples.

### Sequencing

DNA extracts were amplified using primers specific to the V4 variable region of the 16S rRNA gene and sequenced at the University of Connecticut Microbial Analysis Resources and Services center using the standard protocols. Extracts were quantified using the Quant-iT PicoGreen kit (Invitrogen, ThermoFisher Scientific). Partial bacterial 16S rRNA genes (V4, 0.8 picomole each 515F and 806R with Illumina adapters and 8 basepair dual indices [33] were, in triplicate, amplified in 15 ul reactions using GoTaq (Promega) with the addition of 10 mg BSA (New England BioLabs). We added 0.1 femtomole 515F and 806R which do not have the barcodes and adapters to overcome initial primer binding inhibition because the majority of the primers do not match the template priming site. The PCR reaction was incubated at 95°C for 2 minutes, the 30 cycles of 30 s at 95.0°C, 60 s at 50.0 °C, and 60 s at 72.0 °C, followed by a final extension at 72.0 °C for 10 minutes. PCR products were pooled, quantified, and visualized using the QIAxcel DNA Fast Analysis (Qiagen). PCR products were combined using a QIAgility robot for liquid handling after the products were normalized based on the concentration of DNA from 350-420 bp. The pooled PCR products were cleaned using the Mag-Bind RxnPure Plus (Omega Bio-tek) according to the manufacturer’s protocol. The cleaned pool was sequenced on the Illumina MiSeq using v.2 2_250 base-pair kit (Illumina, Inc). Two PCR controls were also sequenced to test for PCR reagent contamination.

### Sequence Quality Control and Processing

Sequence processing for the 181 samples was done using R v.4.1.0 [34]. Sequences were quality controlled, denoised, and merged using *DADA2* v.1.20.0 [35] to create a sample by ASV (amplicon sequence variant) matrix. An ASV is an operational taxonomic unit, defined as any unique sequence that passes stringent quality control. Taxonomy of ASVs was assigned using RDP’s Naïve Bayesian Classifier with the Silva reference database v128 [36,37]. Sequences that were identified as mitochondria, chloroplasts, or that were unable to be confidently assigned to any bacterial phylum were removed. Additionally, the negative controls and the *Decontam* package v.1.12.0 [38] were used to identify and remove likely contaminants. To calculate phylogenetic diversity metrics, a multiple-alignment of all ASVs was performed using the *DECIPHER* package in R [39], and a phylogenetic tree was constructed with the *phangorn* package v.2.4.0 [40].

### Data visualization and statistical analyses

Rarefaction level was determined by generating rarefaction curves using the Shannon diversity index. We rarefied to an even depth (5,464 sequences/sample) and used the rarefied data for all of the following analyses. Alpha diversity was quantified using Shannon diversity index for every sample. We removed samples that did not have sex or age recorded from our dataset before we tested for significant differences in sex and age. The normality of the data was tested using the Shapiro-Wilk test [41]. Alpha diversities were not normally distributed so the Kruskal-Wallis t-test was used for age, sex, state, and flyway at the national level. Data were then subset into the Atlantic (Rhode Island, Connecticut, New York, New Jersey), Central (Kansas and Utah), and Pacific (Nevada, and Washington) flyways and tested for significant differences between age, sex, and state. The Wilcoxon rank-sum t-test [42] with false discovery rate (FDR) *p*-value adjustment method determined what states were significantly different from each other. Taxa less than 1% in each sample were grouped together at each taxonomic rank and “common” microbiota was classified as any taxa present in at least 50% of the samples. Relative abundance data for the phyla, families, and genera were generated for the total dataset as well as the Pacific and Atlantic flyways.

Beta diversity was quantified using three distance metrics: Bray-Curtis (BC), weighted UniFrac (WU), and unweighted UniFrac (UU). Beta diversity was visualized using principal coordinate analysis (PCoA) ordinations. Significance was assessed using permutational multivariate analysis of the variance (PERMANOVA) using the *vegan* command *adonis()* in R [34,43]. Before testing the entire dataset, samples that did not have age or sex recorded were removed (for tests of those specific variables). Data were subset by flyway to test age, sex, and state. The correlation between distance and beta diversity (using Bray-Curtis, weighted UniFrac, and unweighted UniFrac) was calculated with the Mantel test with Spearman’s correlation. Beta diversity distance decay plots were visualized in R and a regression line was generated using a generalized additive model (*gam*) with 95% confidence intervals [44]. As a control, the weighted UniFrac beta diversity distance matrix and the geographic distance matrix were randomized and tested for significance using the Mantel test.

### Supervised learning analysis

The relative abundances at different taxonomic ranks (phylum, class, order, family, and genus) were the features used to predict the state and flyway. Phyla and classes below 1% relative abundance in each sample were grouped into a new category. Orders, families, and genera below 5% in each sample were grouped into a new category. We did not remove the low abundant taxa entirely because the amount of low abundant taxa might have been regional specific. All taxonomic data was reformatted and exported from R. The taxonomic data for each rank was in Jupyter Notebook (python v.3.9.7) [45] and preprocessed the data following a modified version of the pipeline presented by Namkung, 2020 [46]. The primary modification was our use of a Ridge Classifier cross-validation model (Ridge), available from SciKit-learn [47]. We only used locations that had more than 20 samples per state or flyway (Pacific; n = 64, Atlantic; n = 79; WA; n = 41, NV; n = 23, NJ; n = 24, and CT; n = 40). A training and testing ratio was set to 80:20, meaning 80% of the data was used to train the model and the remaining 20% was used to test the model’s accuracy. 10-fold cross-validation was used and the mean accuracy and standard deviation of the cross-validated predictions for each taxonomic rank were recorded. Python code used for these analyses is available at (https://github.com/joshuacgil/Gil_Hird_CAGO_Biogeography_Analysis)

## Results

### Few samples were lost during sequencing and data processing

In total, 182 samples were sequenced and eight samples failed to amplify and/or sequence. After quality control in DADA2, an additional three samples were removed (they had sequence counts of 0 after the non-chimeric sequences were removed). 30,302,294 total reads remained in the final 182 samples. After contaminant, chloroplast, mitochondria, and sequences from unidentified phyla were removed, there were 21,562,720 sequencing reads, at an average of 120,462 reads per sample. Samples were rarefied to an even depth of 5,464 reads per sample; 18 samples were removed for having fewer than 5,000 reads, leaving 161 total samples.

### Canada goose microbiota share large portions of their bacterial microbiota across the United States

17 bacterial phyla were identified that constituted at least 1% of the microbiota in the 161 samples. The most prevalent phyla were *Firmicutes* 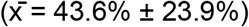, *Proteobacteria* 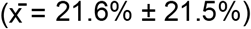, *Actinobacteria*, 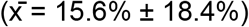, and *Bacteroidetes* 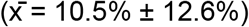. The remaining phyla belonged to *Tenericutes, Fusobacteria, Cyanobacteria, Deferribacteres, Verrucomicrobia*, *Deinococcus*, and *Synergistetes*. *Tenericutes* were found in abundances greater than 5% in 18 samples and ranged from 5.9%-74.1% 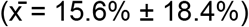. At the state level, the phylum-level composition was largely uniform with *Firmicutes, Proteobacteria, Actinobacteria*, and *Bacteroidetes* in every state and comprising the majority of the phyla (Fig 1). The most abundant phyla in every state were *Firmicutes* and *Proteobacteria. Actinobacteria* and *Bacteroidetes*. Notably, states in the pacific flyway (Washington and Nevada), had much higher relative amounts of *Actinobacteria* (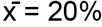 and 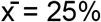 respectively) compared to the other states. The next highest was Connecticut which had 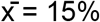 *Actinobacteria*.

**Figure 1:**
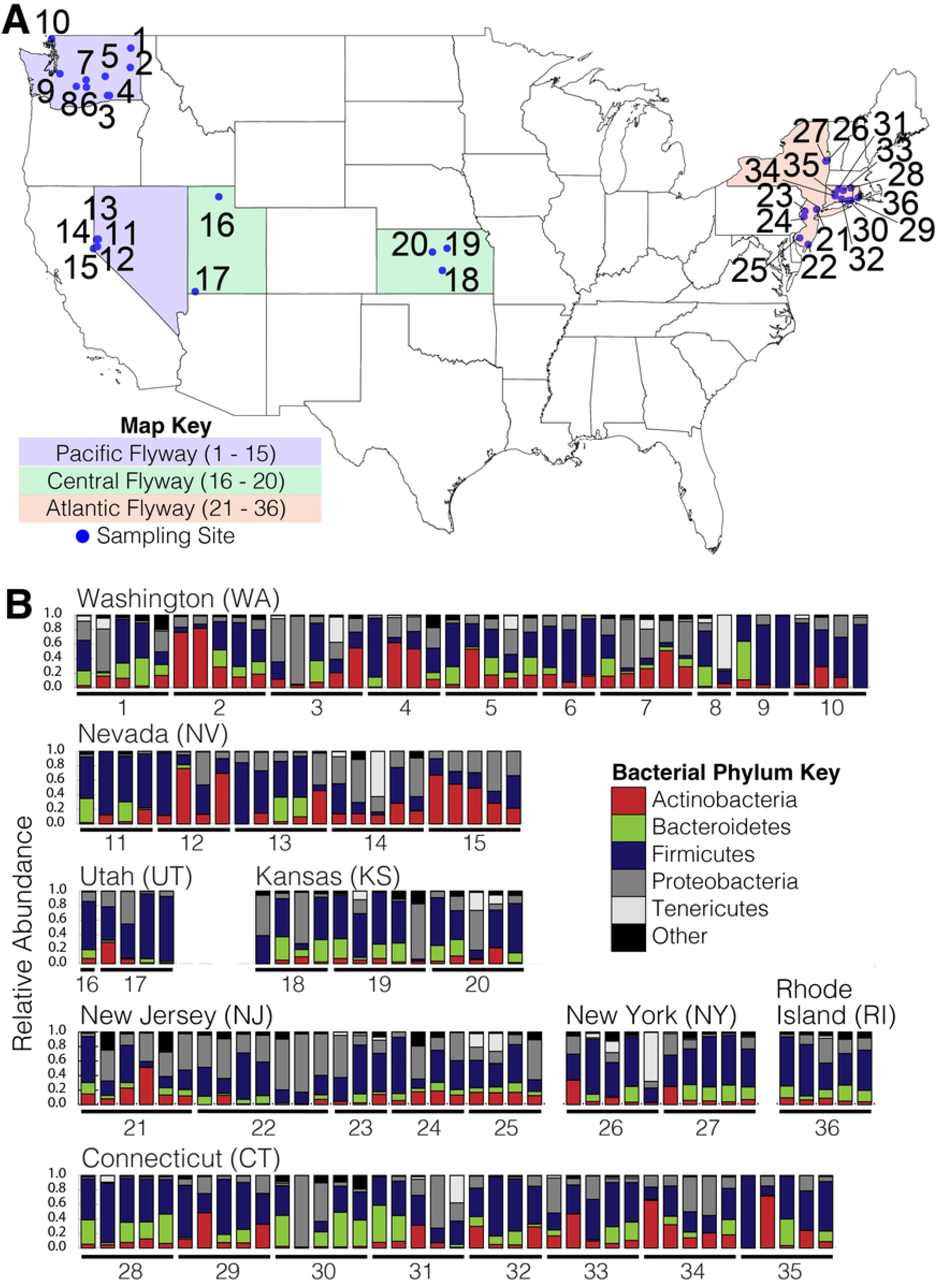
Phylum level relative abundance in Canada goose fecal microbiota. States include Washington (n = 41), Nevada (n = 22), Utah (n = 5), Kansas (n = 14), New Jersey (n = 24), New York (n = 10), Connecticut (n = 40), and Rhode Island (n = 5). Blue dots represent the sampling location. States are colored according to their respective flyways (blue = Pacific, green = Central, and red = Atlantic). “Other” includes phyla <1% of the relative abundance.

Bacteria were considered part of the “common” microbiota if they were present in at least 50% of the samples and comprised at least 1% of the relative abundance. Four phyla were detected across all of the samples (Table 2). The flyways (Pacific and Atlantic) shared the same phyla and the relative abundances of each phylum were similar, except for *Bacteroides* (Table 2). *Bacteroides* was found in 50% of the samples in the Pacific flyway; whereas, it was found in 82% of the samples in the Atlantic flyway samples (Table 2).

**Table 2:**
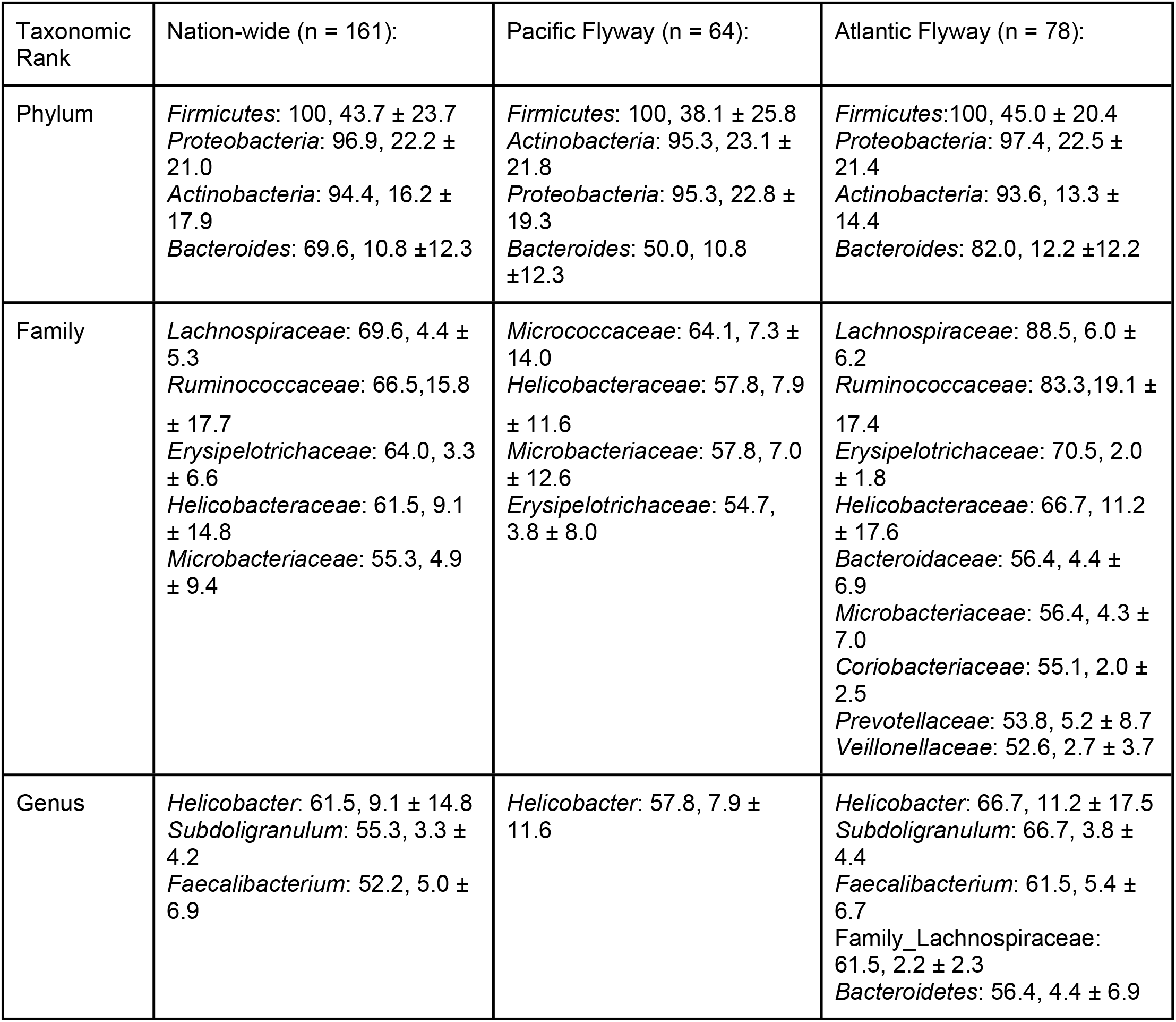
Common taxa (found in >50% of samples) at different taxonomic levels; cells include the taxon, followed by the percentage of samples it was found in, and the average relative abundance ± the standard deviation.

94 families were present in at least one sample at a relative abundance greater than 1%. Five families were present in at least 50% of all samples (*Lachnospiraceae, Ruminococcaceae, Erysipelotrichaceae, Helicobacteraceae*, and *Microbacteriaceae*). The Pacific flyway did not contain *Ruminococcaceae* and *Lachnospiraceae*. The Atlantic flyway had more shared families (9 total), the four additional families were *Bacteroidaceae, Coriobacteriaceae, Prevotellaceae*, and *Veillonellaceae* (Table 2).

185 genera were in at least one sample at a relative abundance greater than 1%. *Helicobacter* was the most abundant genus in all of the samples, as well as within each flyway: *Helicobacter* was found in 66.7% of the Atlantic samples with an average of 11.2% relative abundance and in the Pacific flyway, *Helicobacter* was found in 57.8% of the samples and comprised 7.9% of the relative abundance (Table 2). An ASV from the family *Lachnospiraceae* that was unable to be resolved at the genus level was also a member of the common microbiota. *Faecalibacterium* was found in at least 50% of the samples nationwide and in the Atlantic flyway; however, in the Pacific flyway it was found in only 37.5% of the samples and comprised 3.3% ± 5.3% of the average relative abundance, much lower than the Atlantic flyway. For prevalence and relative abundance of taxa see Table S1 for phyla, Table S2 for families, and Table S3 for genera.

“Common” microbiota was present in at least 50% of the samples and comprised at least 1% of the relative abundance (detection threshold).

### States have significant differences in alpha diversity but flyways do not

There was no difference in alpha diversity of the fecal microbiota of Canada geese between males and females (*p* = 0.73319, X^2^ = 0.3919, df =1), nor between adults and juveniles (*p* = 0.2432, X^2^ = 1.3617, df =1). State of origin was significant (*p* = 0.000298, X^2^ = 3.2518, df = 7, Fig 3). There was no significant difference between the Atlantic, Central, and Pacific flyways (*p =* 0.1967, X^2^ = 3.2518, df = 2). The age and sex within each of the flyways were also not significant (Table S4).

**Figure 2:**
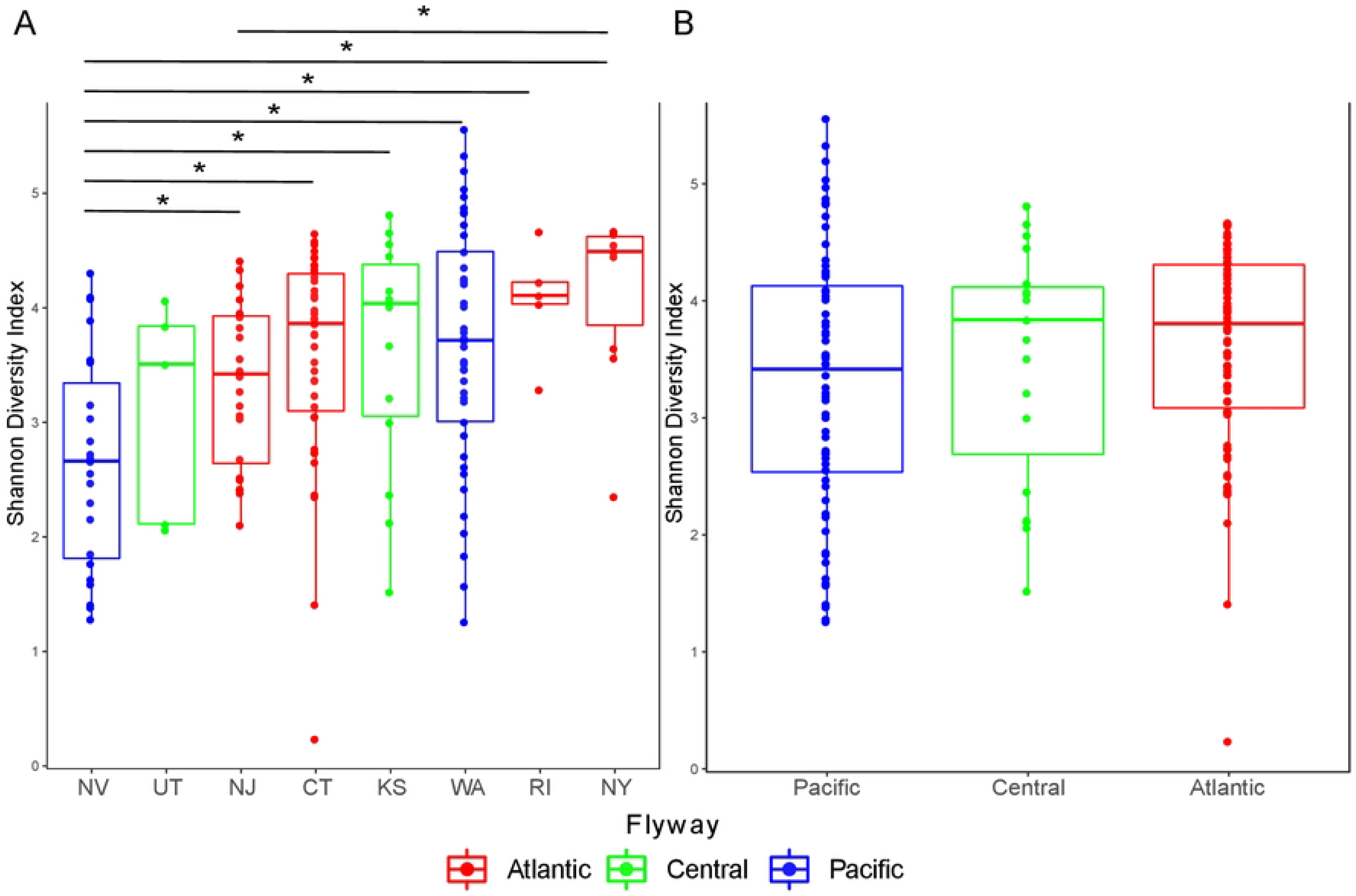
Alpha diversity boxplots for the states and flyways using Shannon diversity index. (A) State box plots include: NV; n = 23, UT; n = 5, NJ; n = 24, CT; n = 40, KS; n = 14, WA; n = 41, RI; n = 5, and NY; n = 10. (B) Flyway box plots include: Pacific; n = 64, Central; n = 19, and Atlantic; n = 79. Significant differences were calculated using the Wilcoxon rank-sum t-test. Significant differences (*p* < 0.05) are denoted by (*). Color corresponds to the flyway (red = Atlantic, green = Central, and blue = Pacific).

**Figure 3:**
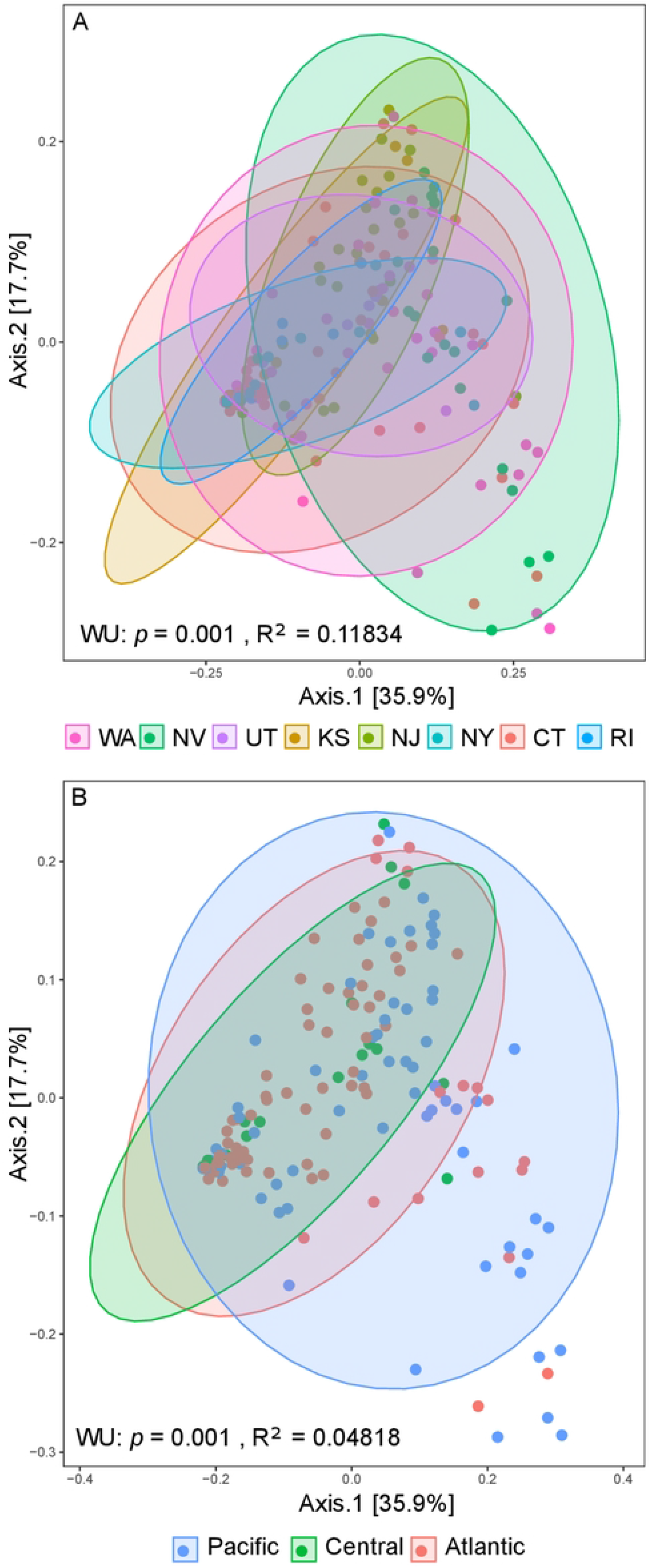
Weighted UniFrac PCoA ordinations. (A) PCoA ordination of the different states, (B) PCoA ordination of the Flyways. PERMANOVA was used to calculate significance and R^2^ values. Ellipses (95% confidence intervals) added around each state and flyway.

Alpha diversity differed significantly (*p* < 0.05) between states. Nevada was significantly lower than every other state except Utah and New Jersey (Wilcoxon rank-sum t-test: NV:CT *p* = 0.0018, NV:KS *p* = 0.0397, NV:NY *p* = 0.0018, NV:RI *p* = 0.0171, and NV:WA *p* = 0.0057). New Jersey and New York were also significantly different (NJ:NY *p* = 0.014); all other pairwise comparisons were not significant (*p* > 0.05). There were no significant differences in alpha diversity between the flyways (Fig 2 and Table S4).

### Beta diversity correlates to state and flyway

Correlations between beta diversity and state, flyway, sex, and age were tested using PERMANOVA. Three distance metrics were used to quantify beta diversity: Bray-Curtis (BC), weighted UniFrac (WU), and unweighted UniFrac (UU). There was a significant correlation in both flyway and state, but the state of origin had larger R^2^ values (Table 3). PCoA ordinations of WU distances showed clustering based on flyway and state (Fig 3). Sex was not significant (Table 3). Age was significant when using BC and UU but both had small R^2^ values and explained very little of the variation observed. The data were then split into their respective flyways and tested again. Sex was not a significant co-factor in any flyway. Age was only significant when using Bray-Curtis in the Pacific flyway. Within the Pacific, Central, and Atlantic flyways, the state of origin was significant when using all three metrics.

**Table 3:**
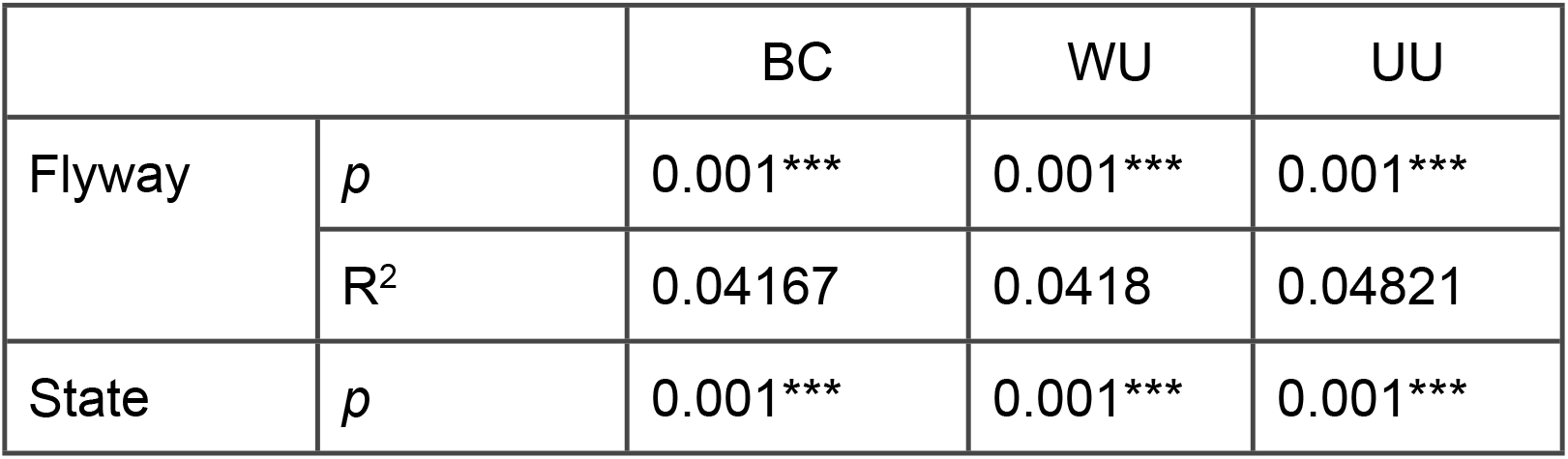
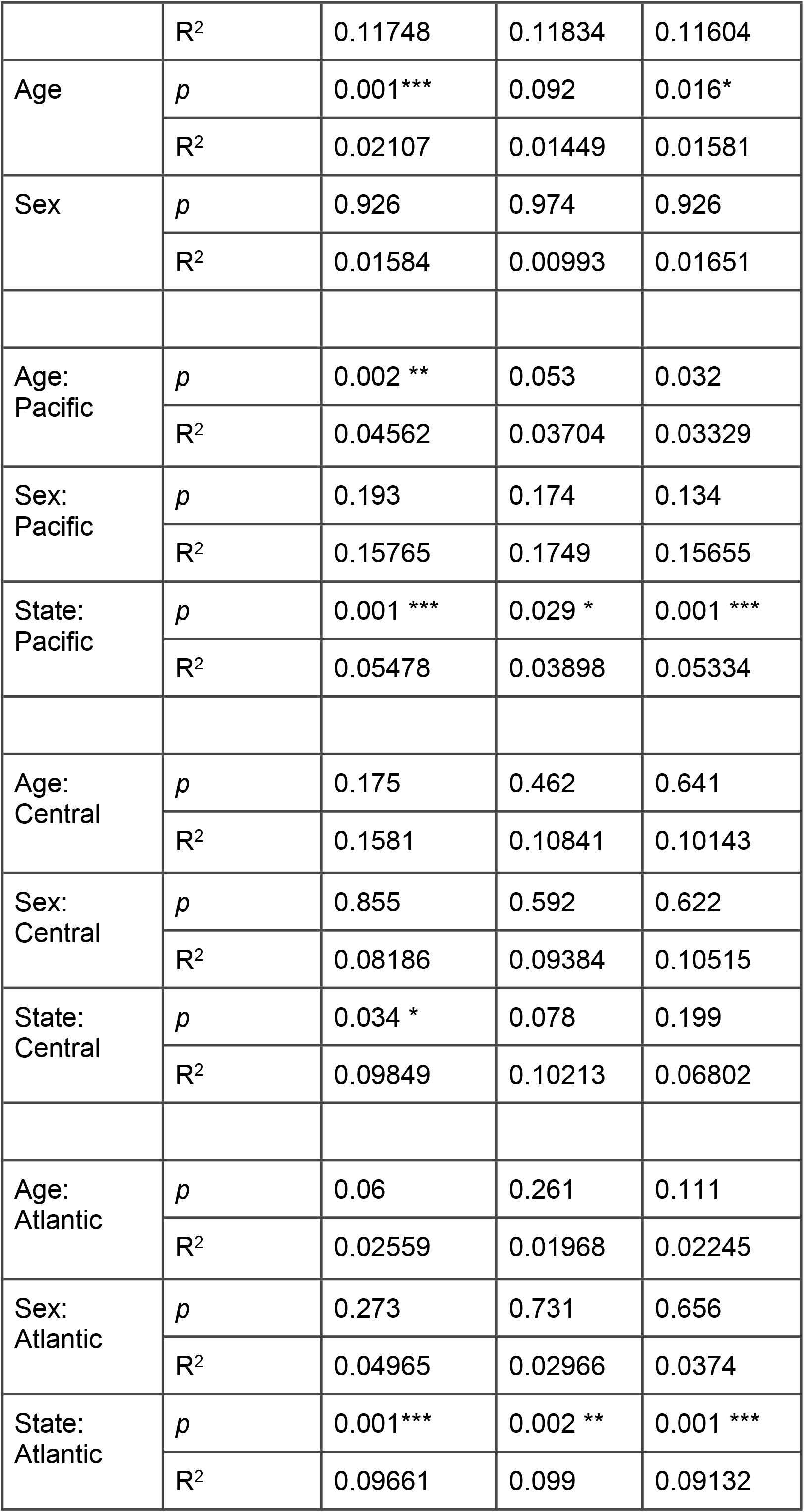
PERMANOVA results (*p*-value and R^2^) testing for correlation of microbiota beta diversity (in Bray-Curtis (BC), weighted UniFrac (WU) and unweighted UniFrac (UU)) for host co-factors (Flyway, State, Age, Sex) on total dataset and after subsetting of the different flyways (e.g., “Age: Pacific” refers to the results of Age in the Pacific flyway). *p < 0.05, **p < 0.01, ***p < 0.001.

|  |  | BC | WU | UU |
| --- | --- | --- | --- | --- |
| Flyway | $p$ | 0.001*** | 0.001*** | 0.001*** |
| | $R^2$ | 0.04167 | 0.0418 | 0.04821 |
| State | $p$ | 0.001*** | 0.001*** | 0.001*** |
|  | R<sup>2</sup> | 0.11748 | 0.11834 | 0.11604 |
| Age | *p* | 0.001*** | 0.092 | 0.016* |
|  | R<sup>2</sup> | 0.02107 | 0.01449 | 0.01581 |
| Sex | *p* | 0.926 | 0.974 | 0.926 |
|  | R<sup>2</sup> | 0.01584 | 0.00993 | 0.01651 |
| Age:<br>Pacific | *p* | 0.002 ** | 0.053 | 0.032 |
|  | R<sup>2</sup> | 0.04562 | 0.03704 | 0.03329 |
| Sex:<br>Pacific | *p* | 0.193 | 0.174 | 0.134 |
|  | R<sup>2</sup> | 0.15765 | 0.1749 | 0.15655 |
| State:<br>Pacific | *p* | 0.001 *** | 0.029 * | 0.001 *** |
|  | R<sup>2</sup> | 0.05478 | 0.03898 | 0.05334 |
| Age:<br>Central | *p* | 0.175 | 0.462 | 0.641 |
|  | R<sup>2</sup> | 0.1581 | 0.10841 | 0.10143 |
| Sex:<br>Central | *p* | 0.855 | 0.592 | 0.622 |
|  | R<sup>2</sup> | 0.08186 | 0.09384 | 0.10515 |
| State:<br>Central | *p* | 0.034 * | 0.078 | 0.199 |
|  | R<sup>2</sup> | 0.09849 | 0.10213 | 0.06802 |
| Age:<br>Atlantic | *p* | 0.06 | 0.261 | 0.111 |
|  | R<sup>2</sup> | 0.02559 | 0.01968 | 0.02245 |
| Sex:<br>Atlantic | *p* | 0.273 | 0.731 | 0.656 |
|  | R<sup>2</sup> | 0.04965 | 0.02966 | 0.0374 |
| State:<br>Atlantic | *p* | 0.001*** | 0.002 ** | 0.001 *** |
|  | R<sup>2</sup> | 0.09661 | 0.099 | 0.09132 |

### Supervised machine learning can accurately predict the origin the fecal samples using taxonomic data

A Ridge model was used to determine whether the state and flyway could be predicted from taxonomic data. The accuracy of a random prediction of the flyway would be 33.3% and the accuracy of a random prediction of the state would be 12.5%. The accuracy of the cross-validated Ridge model for both the Pacific and Atlantic flyways exceeds the random prediction accuracy for every taxonomic rank (Fig 4A). The most accurate predictions for each flyway came from different taxonomic ranks. The Atlantic flyway’s most accurate predictions were derived from genera level data (0.703 ± 0.125); whereas, the Pacific was most accurate at the order level (0.718 ± 0.055).

**Figure 4.**
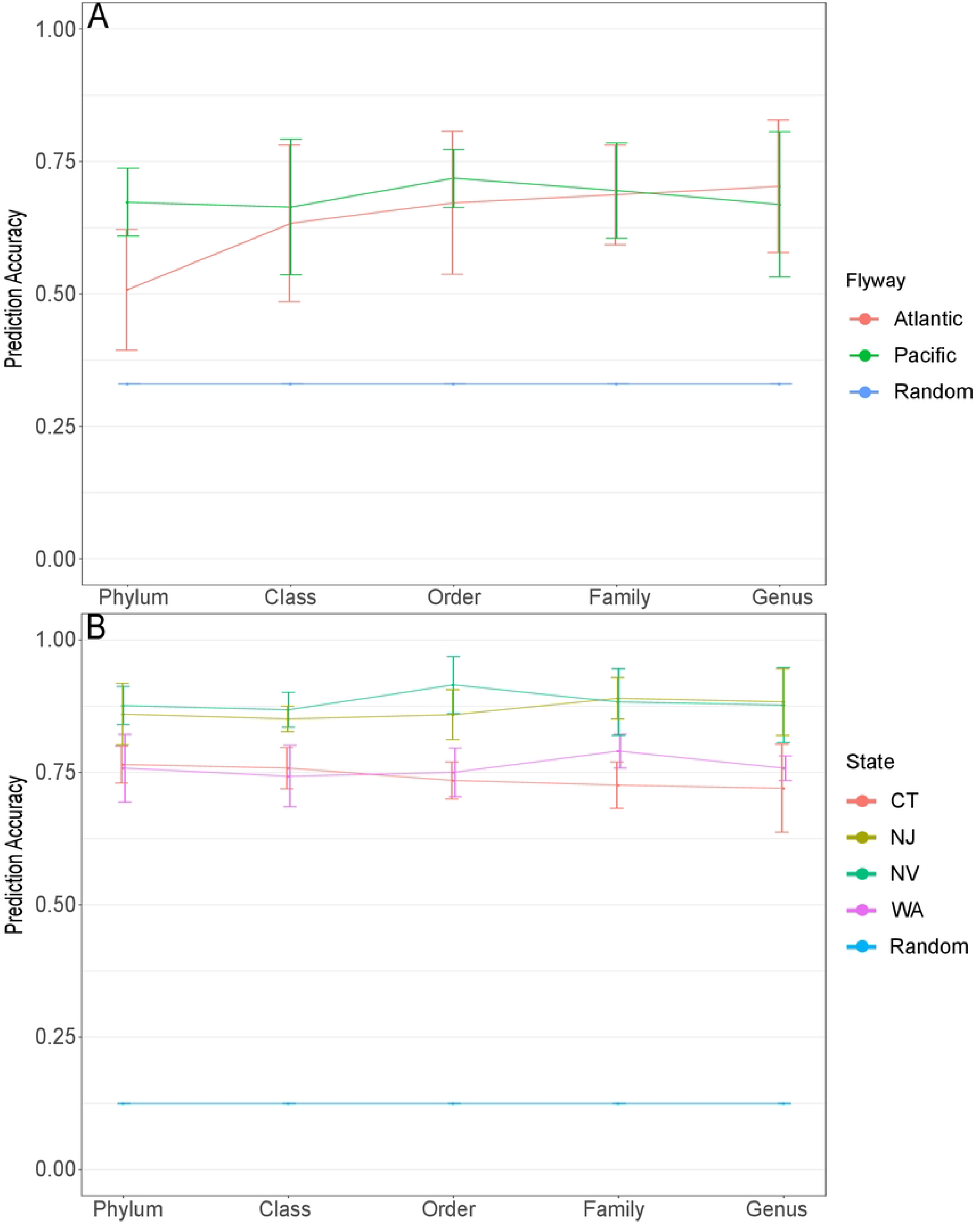
Prediction accuracy of cross-validated Ridge model for each taxonomic rank using locations with >20 samples. Predictions determined using 10-fold cross-validation. (A) Flyway prediction accuracy and theoretical random prediction. (B) State prediction accuracy and theoretical random predictions. Phyla and Classes below 1% relative abundance were grouped into a low abundance category. Orders, families, and genera below 5% relative abundance were grouped into a low abundance category.

The state predictions were better than random predictions regardless of taxonomic rank. The predictions were consistently higher than the flyway predictions with a minimum of 0.72 (CT at genus level). The best predictions for the four states occurred at different taxonomic levels (Fig 4B). The best prediction for CT was at the phylum level (0.765 ± 0.035) and NV was at the order level (0.915 ± 0.054). Both WA and NJ were most accurate at the family level (0.790 ± 0.032 and 0.890 ± 0.039 respectively). See additional file 5 for Ridge model prediction scores.

### Beta diversity positively correlates with geographic distance

Significant and positive correlations between beta diversity and geographic distance were found across the United States (Mantel test: BC; *p* = 0.001, R-stat = 0.1974, WU; *p* = 0.001, R-stat = 0.1141, UU; *p* = 0.001, R-stat 0.1832, Fig 5). Regarding flyway, the Atlantic flyway showed significant correlations between beta diversity and geography when using BC (*p* = 0.003, R-stat = 0.1283) and UU (*p* = 0.016, R-stat = 0.1128), but not WU (*p* = 0.057, R-stat = 0.0585, Fig 5). The Pacific flyway showed greater significant correlations with BC (*p* =0.001, R-stat = 0.2331), WU (*p* = 0.013, R-stat =0.0808), and UU (*p* = 0.001, R-stat = 0.1938, Fig 5)

**Figure 5:**
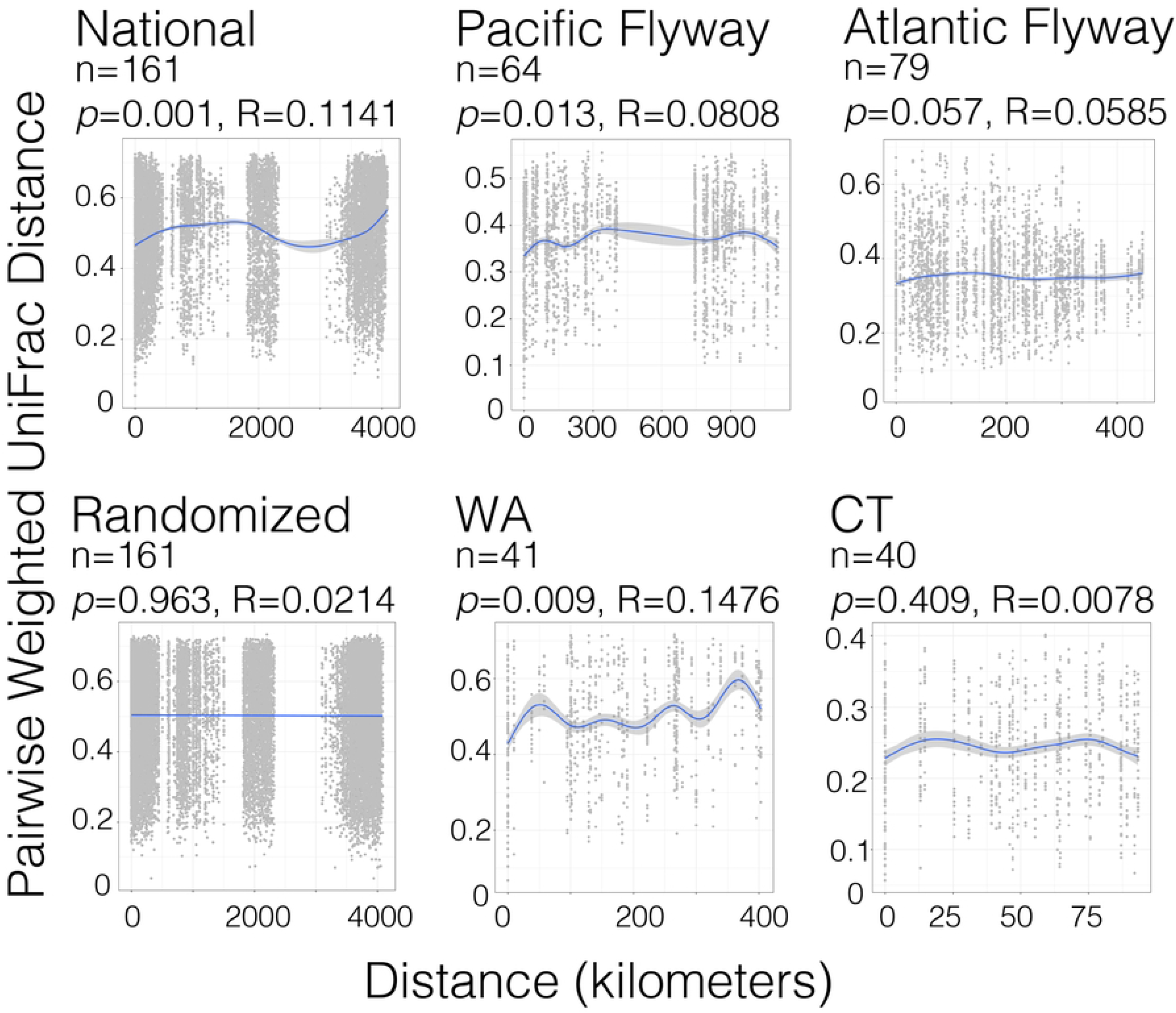
Weighted UniFrac beta diversity distance decay plots in kilometers. Gray dots represent each pairwise comparison between two samples. The regression line in blue with a gray 95% confidence interval was calculated using generalized additive models (*gam*). Significant correlations between distance and beta diversity were tested using the Mantel test, significance (*p* <0.05).

Within the states with the most samples (WA: N = 41 and CT: N = 40), Washington showed significant correlations using all three metrics (BC; *p* =0.001, R-stat = 0.2321, WU; *p* = 0.009, R-stat =0.1476, UU; *p* = 0.002, R-stat = 0.2512, Fig 5). Connecticut had significant correlations with Bray-Curtis (*p* = 0.02, R-stat = 0.1024), but not weighted and unweighted UniFrac (WU; *p* = 0.409, R-stat = 0.0078 and UU; *p* = 0.067, R-stat = 0.0841, Fig 5). To confirm that randomized data did not display significant associations, locations were randomly assigned to pairwise distances and plotted. As expected, randomized data were not significantly associated with distance (*p* = 0.963, R-stat = −0.02139, Fig 5).

## DISCUSSION

The microbiota can confer many benefits to the host, but the factors that shape the microbiota in wild birds, are poorly known [13]. Many microbiota studies use samples from one or several localities but then draw conclusions about the entire species’ microbiota. We aimed to elucidate intraspecies variation in the microbiota and identify the role of geographic distance across a host species’ range on the microbiota. To that end, we characterized and analyzed the fecal microbiota of Canada geese, which have a large range and relatively uniform diet, across ~4000 km in the United States.

Seven of the twenty-eight pairwise comparisons of alpha diversity within states were significant and six of these were comparisons to the state of Nevada. No significant differences were detected when comparing the three different flyways (Pacific, Central, and Atlantic). Alpha diversity was relatively consistent across states and flyways. This suggests that regardless of their environment Canada geese will have comparable bacterial richness and evenness in their microbiota, save a few exceptions (Fig 2). not strongly correlated by location. Further investigation is needed to understand why geese from Nevada might have lower alpha diversity than geese from other states. The differences in diversity might be the result of potential pollution or antibiotic runoff [48–50]. Lower alpha diversity can be associated with active infections [51,52] and the Canada goose population in Nevada may be suffering from an infectious outbreak. More research is required to determine why geese in Nevada have significantly lower alpha diversities; this could represent a temporary health-related issue or perhaps there is another environmental factor that is depressing alpha diversity.

Beta diversity correlated significantly with both state and flyway. The state of origin was a significant co-factor and explained large portions (~11%) of the variation observed (Table 3). Furthermore, all three distance metrics yielded similar results (Table 3). Beta diversity was significantly different depending on the flyway; however, the variation explained by the flyway was much less than what was observed at the state level (~4%, Table 3 and Fig 3). This suggests there are environmental factors that shape the microbiota at more localized levels, and there are fewer or less significant environmental factors at the scale of the flyways.

Supervised machine learning is being increasingly used in microbiology and is often applied to understand the relationship between host-associated microbiota and various disease states [53–55]. We can similarly use supervised machine learning to understand the relationship between the taxonomic composition and locational data. These techniques can be used to predict a sample’s origin based on taxonomic data of the microbiota. Being able to accurately predict the origin of a sample based on the microbiota is additional evidence of a correlation between geography and the microbiota. We would expect purely random predictions of the flyway and state to be 33.3% and 12.5% accurate, respectively. The Ridge model predicted accurate locations for both the states (Range: 72% - 91.5%) and flyways (Range: 50.8% - 71.8%) at rates exceeding the random predictions, regardless of the taxonomic rank (Fig 4). The accuracy of the ML predictions at the different taxonomic ranks shows that the states and flyways are distinct at all taxonomic ranks, despite the general observation that microbiota vary more intra-individually at lower taxonomic ranks (like species or ASVs) and are more stable at higher taxonomic ranks (like phyla). State predictions were more accurate than the flyway predictions (Fig 4). These results are analogous to our beta diversity analysis and suggest that the local environment has a greater effect on the microbiota. A drawback of this analysis is that only two flyways and four states were assessed due to the minimum sample sizes required. (only regions with more than 20 samples were included). Using the Ridge model on regions with less than 20 samples appeared to generate over-fitted models and unreliable results. Further sampling from the Central and Mississippi flyways could be used to verify the relationships observed in the Atlantic and Pacific flyways.

Significant correlations between geography and beta diversity dissimilarity were present at the continental level (Fig 5). Similarly, we saw significant correlations at the flyway level for both the Atlantic and Pacific flyways (Fig 5). The R-stats generated from the Mantel test for the Atlantic flyway were much lower than the Pacific. We looked into WA and CT more in-depth, as they had the largest sample sizes to test if the correlations observed in the flyways will still be present in more localized levels. Fecal samples from Washington had a stronger and more significant relationship between the microbiota and geography than those from Connecticut. This could be explained by the differences in distances between two samples in the different states. The maximum distance between two samples in Connecticut was less than 100 km and the maximum distance between two samples in Washington was ~400 km.

In many microbial systems, the increase in distance correlates to an increase in microbial diversity [17,27,28,56,57]. Canada geese microbiota also adheres to this pattern; however, our data suggest there is a distance threshold that must be breached before significant differences are observable. These data show that Canada geese microbiota are more similar when the geese are geographically closer and more dissimilar when they are more distant from each other. Our results suggest the Canada goose microbiota is influenced predominantly by their local environment; however, we would like to note that host genetics might also be autocorrelated with geography and confounding the geographic signals. There are seven distinct subspecies large bodied Canada geese that are region specific [58,59], but little is known about the genetic diversity of the resident populations across North America. The different resident populations in the different regions are likely not mixing with populations from other flyways and it is probable these allopatric populations are undergoing some level of genetic diversification. This genetic diversification could lead to region specific selective pressures on the microbiota that are technically derived from the host not the local environment. Further research on the genetic diversity of resident Canada goose populations across the United States could be used to determine how their phylogeny correlates with the microbiota diversity. These experiments would reveal if resident Canada goose populations are undergoing phylosymbiosis [60].

The majority of the samples had the following phyla: *Firmicutes* (100%), *Proteobacteria* (96%), *Bacteroidetes* (94%), and *Actinobacteria* (69.6%) (Fig 1 and Table 2). Five families and three genera were also found to be a part of the “common” microbiota, where “common” is defined as taxa found in at least 50% of the samples. Samples within the Atlantic flyway shared more families (nine) and genera (five) than those within the Pacific flyway, which had four families and one genus (Table 2). Large amounts of the genera *Faecalibacterium* and *Subdoligranulum* were detected in the majority of our samples. Both genera are obligate anaerobes and are associated with fermentation reactions that produce short-chain fatty acids [61–63]. These fatty acids are the byproduct of microbial fermentation of plant matter and can then be used by the host [8,64–67]. Because these genera are in most of our samples, it is possible that they are key constituents of the microbiota and are crucial for the goose’s digestion of highly fibrous food. This consistency of these microbes in Canada geese across North America raises an interesting question. How do these geese acquire an obligate anaerobe as part of their microbiota across such a vast range? It is unlikely they acquired it from the environment through horizontal transmission as the external environment these geese inhabit is obviously oxygenated. We suspect there is some form of vertical transmission, oviposition is a mode of vertical transmission from parent to offspring [68], but since these microbes are anaerobes, they would not survive on the eggshell long enough for the goslings to become exposed. Vertical transmission *in ovo* has been documented in other oviparous animals [69], we suspect this is the mode of transmission for these microbes. More research is required to verify this and identify what other members of the microbiota are transmitted *in ovo*. The high abundance and prevalence of *Helicobacter* may be a matter of human concern since many *Helicobacter* species are potential pathogens [70]. Other potential zoonotic genera were also present in fewer samples and abundances: *Campylobacter* (42/161 samples), *Escherichia* (16/161 samples), and *Mycobacterium* (14/161 samples) (Table S3). Although Canada geese have not been a major concern for public health, these findings indicate geese may be a potential vector of zoonotic pathogens threatening humans, other animals and crops [71–75].

Intrinsic factors like age and sex have been significant co-factors in microbiota composition in many host organisms [18,76–78]; however, sex does not seem to be a major co-factor for Canada geese in either alpha or beta diversity. Age was a significant co-factor for beta diversity using two distance metrics in the Pacific flyway. We hypothesize this is because the samples collected from Nevada were all adults; whereas, the samples from Washington were a mix of adults and juveniles, which would confound the age results with the geographic signal. If we had juvenile samples from Nevada, there may have been no significant differences connected to age. Sex was an insignificant co-factor and had no detectable influence on either alpha or beta diversity in Canada geese. These results corroborate existing work that showed age and sex had little bearing on either alpha or beta diversity in Canada geese [79]. Conclusion:

The microbiota of Canada geese are influenced more by geography than other co-factors. Geese that are closer together have more similar microbiota and geese farther apart have more dissimilar microbiota. Samples from within the same state were more similar than samples from within the same flyway, but both explained a significant amount of microbiota variation. This suggests that there are environmental pressures affecting the microbiota at local levels more than the continental level. There are significant differences in the microbiota across the species range, but there are still strong similarities resulting in conserved microbial taxa in the majority of the geese sampled. This is likely because geese are folivores and have similar diets regardless of their location, likely resulting in a convergence in microbial taxa that are specialized in fiber degradation. Our results also show that intrinsic factors like age and sex are not significant factors affecting the microbiota and the environment is the more critical component of microbiota assembly.

## Acknowledgements

We would like to thank Dr. Elizabeth Herder, Ryan Duggan, and Todd Peters for helping collect samples from Connecticut. We would like to thank our collaborators from across the United States who either collected fecal samples or allowed us to come and collect samples during their routine banding: Kyle Spragens and Matthew Wilson with Washington Department of Fish & Wildlife, Russel Woolstenhulme with Nevada Department of Wildlife, Blair Stringham with Utah Division of Wildlife Resources, Tom Bidrowski and Andrew Friesen Kansas Department of Wildlife, Ted Nichols with New Jersey Division of Fish and Wildlife, Thomas Bell and Joshua Stiller with New York Department of Environmental Conservation, Kelly Kubik and Min Huang with Connecticut’s Department of Energy and Environmental Protection, and Jennifer Kilburn with Rhode Island Department of Environmental Management.

## Ethics Statement

All samples were collected passively while Canada geese were undergoing routine banding performed by State Fish and Wildlife Departments.

## Supporting Information Files

File S1 Metadata file used for analyses

File S2 Table S1 Phylum Level relative abundances

File S3 Table S2 Family Level relative abundances

File S4 Table S3 Genus level relative abundances

File S5 Table S4 Alpha diversity statistics

File S6 Table S6 Machine learning prediction scores.

## Data Availability

All sequences are in the process of being upload into the Short Read Archive (SRA). Scripts used to generate the results can be found in the following GitHub repository https://github.com/joshuacgil/Gil_Hird_CAGO_Biogeography_Analysis

## REFERENCES

1. McFall-Ngai M, Hadfield MG, Bosch TCG, Carey HV, Domazet-Lošo T, Douglas AE, et al. Animals in a bacterial world, a new imperative for the life sciences. Proc Natl Acad Sci U S A. 2013;110: 3229–3236.

2. Round JL, Mazmanian SK. The gut microbiota shapes intestinal immune responses during health and disease. Nat Rev Immunol. 2009;9: 313–323.

3. Schippa S, Conte MP. Dysbiotic events in gut microbiota: impact on human health. Nutrients. 2014;6: 5786–5805.

4. Kim CH. Immune regulation by microbiome metabolites. Immunology. 2018;154: 220–229.

5. Patterson JA, Burkholder KM. Application of prebiotics and probiotics in poultry production. Poult Sci. 2003;82: 627–631.

6. Bucher MG, Zwirzitz B, Oladeinde A, Cook K, Plymel C, Zock G, et al. Reused poultry litter microbiome with competitive exclusion potential against Salmonella Heidelberg. J Environ Qual. 2020;49: 869–881.

7. Hanning I, Diaz-Sanchez S. The functionality of the gastrointestinal microbiome in non-human animals. Microbiome. 2015;3: 51.

8. Remington TE. Why do grouse have ceca? A test of the fiber digestion theory. J Exp Zool Suppl. 1989;3: 87–94.

9. Garcia DM. The role of the giant Canada goose (Branta canadensis maxima) cecum in nutrition. 2006. Available: https://mospace.umsystem.edu/xmlui/handle/10355/4354

10. Liu G, Luo X, Zhao X, Zhang A, Jiang N, Yang L, et al. Gut microbiota correlates with fiber and apparent nutrients digestion in goose. Poult Sci. 2018;97: 3899–3909.

11. Bodawatta KH, Hird SM, Grond K, Poulsen M, Jønsson KA. Avian gut microbiomes taking flight. Trends Microbiol. 2021. doi:10.1016/j.tim.2021.07.003

12. Colston TJ, Jackson CR. Microbiome evolution along divergent branches of the vertebrate tree of life: what is known and unknown. Mol Ecol. 2016;25: 3776–3800.

13. Waite DW, Taylor MW. Exploring the avian gut microbiota: current trends and future directions. Front Microbiol. 2015;6: 673.

14. Song SJ, Sanders JG, Delsuc F, Metcalf J, Amato K, Taylor MW, et al. Comparative Analyses of Vertebrate Gut Microbiomes Reveal Convergence between Birds and Bats. MBio. 2020;11. doi:10.1128/mBio.02901-19

15. Grond K, Sandercock BK, Jumpponen A, Zeglin LH. The avian gut microbiota: community, physiology and function in wild birds. J Avian Biol. 2018;49: e01788.

16. Capunitan DC, Johnson O, Terrill RS, Hird SM. Evolutionary signal in the gut microbiomes of 74 bird species from Equatorial Guinea. Mol Ecol. 2020;29: 829–847.

17. Fleischer R, Risely A, Hoeck PEA, Keller LF, Sommer S. Mechanisms governing avian phylosymbiosis: Genetic dissimilarity based on neutral and MHC regions exhibits little relationship with gut microbiome distributions of Galápagos mockingbirds. Ecol Evol. 2020;10: 13345–13354.

18. Ballou AL, Ali RA, Mendoza MA, Ellis JC, Hassan HM, Croom WJ, et al. Development of the Chick Microbiome: How Early Exposure Influences Future Microbial Diversity. Front Vet Sci. 2016;3: 2.

19. Hird SM, Carstens BC, Cardiff SW, Dittmann DL, Brumfield RT. Sampling locality is more detectable than taxonomy or ecology in the gut microbiota of the brood-parasitic Brown-headed Cowbird (Molothrus ater). PeerJ. 2014;2: e321.

20. Hird SM, Sánchez C, Carstens BC, Brumfield RT. Comparative Gut Microbiota of 59 Neotropical Bird Species. Front Microbiol. 2015;6: 1403.

21. Kreisinger J, Kropáčková L, Petrželková A, Adámková M, Tomášek O, Martin J-F, et al. Temporal Stability and the Effect of Transgenerational Transfer on Fecal Microbiota Structure in a Long Distance Migratory Bird. Front Microbiol. 2017;8: 50.

22. Waite DW, Taylor MW. Characterizing the avian gut microbiota: membership, driving influences, and potential function. Front Microbiol. 2014;5: 223.

23. Mahmood T, Guo Y. Dietary fiber and chicken microbiome interaction: Where will it lead to? Anim Nutr. 2020;6: 1–8.

24. Herder EA, Spence AR, Tingley MW, Hird SM. Elevation Correlates With Significant Changes in Relative Abundance in Hummingbird Fecal Microbiota, but Composition Changes Little. Frontiers in Ecology and Evolution. 2021;8: 534.

25. Pearce DS, Hoover BA, Jennings S, Nevitt GA, Docherty KM. Morphological and genetic factors shape the microbiome of a seabird species (Oceanodroma leucorhoa) more than environmental and social factors. Microbiome. 2017;5: 146.

26. Trevelline BK, Sosa J, Hartup BK, Kohl KD. A bird’s-eye view of phylosymbiosis: weak signatures of phylosymbiosis among all 15 species of cranes. Proc Biol Sci. 2020;287: 20192988.

27. King AJ, Freeman KR, McCormick KF, Lynch RC, Lozupone C, Knight R, et al. Biogeography and habitat modelling of high-alpine bacteria. Nat Commun. 2010;1: 53.

28. Angermeyer A, Crosby SC, Huber JA. Salt marsh sediment bacterial communities maintain original population structure after transplantation across a latitudinal gradient. PeerJ. 2018;6: e4735.

29. Lu J, Santo Domingo JW, Hill S, Edge TA. Microbial diversity and host-specific sequences of Canada goose feces. Appl Environ Microbiol. 2009;75: 5919–5926.

30. Gorham TJ, Lee J. Pathogen Loading From Canada Geese Faeces in Freshwater: Potential Risks to Human Health Through Recreational Water Exposure. Zoonoses Public Health. 2016;63: 177–190.

31. Marie-Christine Cadieux, Gauthier G, Hughes RJ. Feeding Ecology of Canada Geese (Branta canadensis interior) in Sub-Arctic Inland Tundra during Brood-Rearing (Écologie alimentaire de Branta canadensis interior pendant la période d’élevage des jeunes dans un milieu d’eau douce sub-arctique). Auk. 2005;122: 144–157.

32. Buchsbaum R, Wilson J, Valiela I. Digestibility of plant constitutents by Canada geese and Atlantic Brant. Ecology. 1986;67: 386–393.

33. Kozich JJ, Westcott SL, Baxter NT, Highlander SK, Schloss PD. Development of a dual-index sequencing strategy and curation pipeline for analyzing amplicon sequence data on the MiSeq Illumina sequencing platform. Appl Environ Microbiol. 2013;79: 5112–5120.

34. Development Core Team RR. R: A language and environment for statistical computing. 2019.

35. Callahan BJ, McMurdie PJ, Rosen MJ, Han AW, Johnson AJA, Holmes SP. DADA2: High-resolution sample inference from Illumina amplicon data. Nat Methods. 2016;13: 581–583.

36. Wang Q, Garrity GM, Tiedje JM, Cole JR. Naive Bayesian classifier for rapid assignment of rRNA sequences into the new bacterial taxonomy. Appl Environ Microbiol. 2007;73: 5261–5267.

37. Quast C, Pruesse E, Yilmaz P, Gerken J, Schweer T, Yarza P, et al. The SILVA ribosomal RNA gene database project: improved data processing and web-based tools. Nucleic Acids Res. 2013;41: D590–6.

38. Davis NM, Proctor DM, Holmes SP, Relman DA, Callahan BJ. Simple statistical identification and removal of contaminant sequences in marker-gene and metagenomics data. Microbiome. 2018;6: 226.

39. Wright ES. DECIPHER: harnessing local sequence context to improve protein multiple sequence alignment. BMC Bioinformatics. 2015;16: 322.

40. Schliep KP. phangorn: phylogenetic analysis in R. Bioinformatics. 2011;27: 592–593.

41. Shapiro SS, Wilk MB. An analysis of variance test for normality (complete samples). Biometrika. 1965;52: 591–611.

42. Wilcoxon F. Individual comparisons by ranking methods. Biom. Bull., 1, 80–83. 1945.

43. Oksanen J, Kindt R, Legendre P, O’Hara B, Stevens MHH, Oksanen MJ, et al. The vegan package. Community ecology package. 2007;10: 631–637.

44. Hastie T, Tibshirani R. Generalized Additive Models. SSO Schweiz Monatsschr Zahnheilkd. 1986;1: 297–310.

45. Van Rossum G, Drake FL. Python 3 Reference Manual: (Python Documentation Manual Part 2). CreateSpace Independent Publishing Platform; 2009.

46. Namkung J. Machine learning methods for microbiome studies. J Microbiol. 2020;58: 206–216.

47. Pedregosa, Varoquaux, Gramfort, Michel, Thirion, Grisel, et al. Scikit-learn: Machine Learning in Python. J Mach Learn Res. 2011;12: 2825–2830.

48. Kraemer SA, Ramachandran A, Perron GG. Antibiotic Pollution in the Environment: From Microbial Ecology to Public Policy. Microorganisms. 2019;7. doi:10.3390/microorganisms7060180

49. Zhu W, Yang D, Chang L, Zhang M, Zhu L, Jiang J. Animal gut microbiome mediates the effects of antibiotic pollution on an artificial freshwater system. J Hazard Mater. 2022;425: 127968.

50. Xue X, Jia J, Yue X, Guan Y, Zhu L, Wang Z. River contamination shapes the microbiome and antibiotic resistance in sharpbelly (Hemiculter leucisculus). Environ Pollut. 2021;268: 115796.

51. Ganz HH, Doroud L, Firl AJ, Hird SM, Eisen JA, Boyce WM. Community-Level Differences in the Microbiome of Healthy Wild Mallards and Those Infected by Influenza A Viruses. mSystems. 2017;2. doi:10.1128/mSystems.00188-16

52. Hird SM, Ganz H, Eisen JA, Boyce WM. The Cloacal Microbiome of Five Wild Duck Species Varies by Species and Influenza A Virus Infection Status. mSphere. 2018;3. doi:10.1128/mSphere.00382-18

53. Marcos-Zambrano LJ, Karaduzovic-Hadziabdic K, Loncar Turukalo T, Przymus P, Trajkovik V, Aasmets O, et al. Applications of Machine Learning in Human Microbiome Studies: A Review on Feature Selection, Biomarker Identification, Disease Prediction and Treatment. Front Microbiol. 2021;12: 634511.

54. Thompson J, Johansen R, Dunbar J, Munsky B. Machine learning to predict microbial community functions: An analysis of dissolved organic carbon from litter decomposition. bioRxiv. 2019. p. 599704. doi:10.1101/599704

55. Liu Y-X, Qin Y, Chen T, Lu M, Qian X, Guo X, et al. A practical guide to amplicon and metagenomic analysis of microbiome data. Protein Cell. 2021;12: 315–330.

56. Astorga A, Oksanen J, Luoto M, Soininen J, Virtanen R, Muotka T. Distance decay of similarity in freshwater communities: do macro-and microorganisms follow the same rules?: Decay of similarity in freshwater communities. Glob Ecol Biogeogr. 2012;21: 365–375.

57. Bueno de Mesquita CP, Nichols LM, Gebert MJ, Vanderburgh C, Bocksberger G, Lester JD, et al. Structure of Chimpanzee Gut Microbiomes across Tropical Africa. mSystems. 2021;6: e0126920.

58. Shields GF, Wilson AC. Subspecies of the Canada Goose (Branta canadensis) Have Distinct Mitochondrial DNA’s. Evolution. 1987;41: 662–666.

59. Scribner KT, Talbot SL, Pearce JM, Pierson BJ, Bollinger KS, Derksen DV. Phylogeography of Canada Geese (Branta Canadensis) in Western North America. Auk. 2003;120: 889–907.

60. Kohl KD. Ecological and evolutionary mechanisms underlying patterns of phylosymbiosis in host-associated microbial communities. Philos Trans R Soc Lond B Biol Sci. 2020;375: 20190251.

61. Van Hul M, Le Roy T, Prifti E, Dao MC, Paquot A, Zucker J-D, et al. From correlation to causality: the case of Subdoligranulum. Gut Microbes. 2020;12: 1–13.

62. Yang H, Xiao Y, Gui G, Li J, Wang J, Li D. Microbial community and short-chain fatty acid profile in gastrointestinal tract of goose. Poult Sci. 2018;97: 1420–1428.

63. Holmstrøm K, Collins MD, Møller T, Falsen E, Lawson PA. Subdoligranulum variabile gen. nov., sp. nov. from human feces. Anaerobe. 2004;10: 197–203.

64. He J, Zhang P, Shen L, Niu L, Tan Y, Chen L, et al. Short-Chain Fatty Acids and Their Association with Signalling Pathways in Inflammation, Glucose and Lipid Metabolism. Int J Mol Sci. 2020;21. doi:10.3390/ijms21176356

65. Macfarlane S, Macfarlane GT. Regulation of short-chain fatty acid production. Proc Nutr Soc. 2003;62: 67–72.

66. Flint HJ, Scott KP, Louis P, Duncan SH. The role of the gut microbiota in nutrition and health. Nat Rev Gastroenterol Hepatol. 2012;9: 577–589.

67. Newbold CJ, Ramos-Morales E. Review: Ruminal microbiome and microbial metabolome: effects of diet and ruminant host. Animal. 2020;14: s78–s86.

68. Bunker ME, Elliott G, Heyer-Gray H, Martin MO, Arnold AE, Weiss SL. Vertically transmitted microbiome protects eggs from fungal infection and egg failure. Anim Microbiome. 2021;3: 43.

69. Trevelline BK, MacLeod KJ, Knutie SA, Langkilde T, Kohl KD. In ovo microbial communities: a potential mechanism for the initial acquisition of gut microbiota among oviparous birds and lizards. Biol Lett. 2018;14. doi:10.1098/rsbl.2018.0225

70. Green HC, Dick LK, Gilpin B, Samadpour M, Field KG. Genetic markers for rapid PCR-based identification of gull, Canada goose, duck, and chicken fecal contamination in water. Appl Environ Microbiol. 2012;78: 503–510.

71. Altekruse SF, Stern NJ, Fields PI, Swerdlow DL. Campylobacter jejuni--an emerging foodborne pathogen. Emerg Infect Dis. 1999;5: 28–35.

72. Friedman CR. Epidemiology of Campylobacter jejuni infections on the united states and other industrialized nations. Campylobacter. 2000; 121–138.

73. Graczyk TK, Fayer R, Trout JM, Lewis EJ, Farley CA, Sulaiman I, et al. Giardia sp. cysts and infectious Cryptosporidium parvum oocysts in the feces of migratory Canada geese (Branta canadensis). Appl Environ Microbiol. 1998;64: 2736–2738.

74. Middleton JH, Ambrose A. Enumeration and antibiotic resistance patterns of fecal indicator organisms isolated from migratory Canada geese (Branta canadensis). J Wildl Dis. 2005;41: 334–341.

75. Gil JC, Hird SM. Multi-omics characterization of the Canada goose fecal microbiome reveals selective efficacy of simulated metagenomes. Research Square. 2022. doi:10.21203/rs.3.rs-1498492/v1

76. Reveles KR, Patel S, Forney L, Ross CN. Age-related changes in the marmoset gut microbiome. Am J Primatol. 2019;81: e22960.

77. Janiak MC, Montague MJ, Villamil CI, Stock MK, Trujillo AE, DePasquale AN, et al. Age and sex-associated variation in the multi-site microbiome of an entire social group of free-ranging rhesus macaques. Microbiome. 2021;9: 68.

78. Couper LI, Kwan JY, Ma J, Swei A. Drivers and patterns of microbial community assembly in a Lyme disease vector. Ecol Evol. 2019;9: 7768–7779.

79. Drovetski SV, O’Mahoney M, Ransome EJ, Matterson KO, Lim HC, Chesser RT, et al. Spatial Organization of the Gastrointestinal Microbiota in Urban Canada Geese. Sci Rep. 2018;8: 3713.

